# Chlorpromazine inhibits EAG1 channels by altering the coupling between the PAS, CNBH and pore domains

**DOI:** 10.1101/2024.02.23.581826

**Authors:** Mahdi Ghorbani, Ze-Jun Wang, Xi Chen, Purushottam B. Tiwari, Jeffery B. Klauda, Tinatin I. Brelidze

## Abstract

EAG1 depolarization-activated potassium selective channels are important targets for treatment of cancer and neurological disorders. EAG1 channels are formed by a tetrameric subunit assembly with each subunit containing an N-terminal Per-Arnt-Sim (PAS) domain and C-terminal cyclic nucleotide-binding homology (CNBH) domain. The PAS and CNBH domains from adjacent subunits interact and form an intracellular tetrameric ring that regulates the EAG1 channel gating, including the movement of the voltage sensor domain (VSD) from closed to open states. Small molecule ligands can inhibit EAG1 channels by binding to their PAS domains. However, the allosteric pathways of this inhibition are not known. Here we show that chlorpromazine, a PAS domain small molecule binder, alters interactions between the PAS and CNBH domains and decreases the coupling between the intracellular tetrameric ring and the pore of the channel, while having little effect on the coupling between the PAS and VSD domains. In addition, chlorpromazine binding to the PAS domain did not alter Cole-Moore shift characteristic of EAG1 channels, further indicating that chlorpromazine has no effect on VSD movement from the deep closed to opened states. Our study provides a framework for understanding global pathways of EAG1 channel regulation by small molecule PAS domain binders.

## INTRODUCTION

Voltage-gated potassium channels (VGPC) are essential for cellular signaling as they are key players in controlling membrane potential [1;2]. VGPC share a common architecture, including a tetrameric assembly from subunits containing six membrane-spanning segments, with segments one through four (S1-S4) forming the voltage sensor domain (VSD) and segments five and six (S5-S6) forming the centrally located pore domain (PD) [1;2]. While the general architecture is similar for VGPC, their distinct functional features are frequently conferred by intracellular domains. The intracellular domains of VGPC function as regulatory and ligand-binding modules, and are promising targets for pharmacological regulation and drug design [3]. Mechanisms of VGPC regulation by ligand binding to their intracellular domains are not well understood.

Ether-a-go-go (EAG1) VGPC, also known as KCNH1 and Kv10.1, is a founding member of the KCNH channel family of ion channels and a key regulator of neuronal excitability [4–9]. While under normal conditions, EAG1 channels are predominantly expressed in the brain, they are overexpressed in the majority of primary human tumors and inhibition of their activity decreases tumor progression [10–16]. EAG1 channels contain the intracellular N-terminal Per-Arnt-Sim (PAS) domains that interact with the intracellular C-terminal cyclic nucleotide-binding homology (CNBH) domains from adjacent subunits, forming an intracellular tetrameric PAS/CNBH domain ring located at the entrance to the pore of the channel [17;18]. Through the interactions with the CNBH domains and also interactions with VSD, PAS domains regulate EAG1 channel gating, including a delayed opening from extreme hyperpolarized potentials (Cole-Moore shift) [17–23]. Recent studies show that the PAS domains of EAG1 channels can also function as ligand-binding domains. Small molecule ligands, such as tricyclic drugs chlorpromazine (CPZ) and imipramine, and undecylenic acid directly bind to the PAS domains and inhibit EAG1 currents through this binding [24–26]. How binding of the small molecule ligands to the intracellularly located PAS domains can affect current flow through the centrally located pore of the channel is not known. Although, KCNH channels are the only ion channels containing PAS domains, the PAS domain fold is quite common in other proteins and is frequently targeted for drug design [27–30]. It is highly likely that many more EAG1 PAS domain small molecule binders will be identified in the future. Therefore, uncovering the mechanisms of EAG1 channel regulation by PAS domain small molecule binders is important for understanding the fundamental steps in the channel gating and for development of novel therapeutics targeting EAG1 channels.

Here we used a combination of molecular dynamics (MD) simulations, surface plasmon resonance (SPR) technique, and network analysis to show that CPZ inhibits EAG1 channels by altering the interactions between the PAS and CNBH domains and decreasing the coupling between the intracellular PAS/CNBH domain tetrameric ring and channel pore domain. In addition, CPZ had no effect on the Cole-Moore shift in EAG1 channels, indicating that it does not affect VSD movement from deep hyperpolarized closed states to the open state. The uncovered allosteric pathways of EAG1 channel inhibition by CPZ provides an initial framework for understanding ligand-dependent gating in EAG1 and other KCNH channels.

### CPZ alters interactions between the PAS and CNBH domains in EAG1 channels

To gain insight into the structural mechanism of EAG1 current inhibition due to CPZ binding to the PAS domain, we performed MD simulations of full-length EAG1 (EAG1) channels in the absence and presence of CPZ bound to the PAS domains of all four subunits. The mEAG1 model used for the MD simulations was built using the cryo-EM structure of full-length rEAG1 channels (pdb id: 5K7L) [17] stripped of calmodulin and embedded into a lipid membrane (Fig. 1a). The generated model contained the N-terminal PAS domain, VSD and PD, and the C-linker connecting the C-terminal CNBH domain to the S6 segment of PD (Fig. 1a). MD simulations were run at room temperature for 1 μs for both CPZ-unbound (apo) and CPZ-bound EAG1 models.

**Fig 1.**
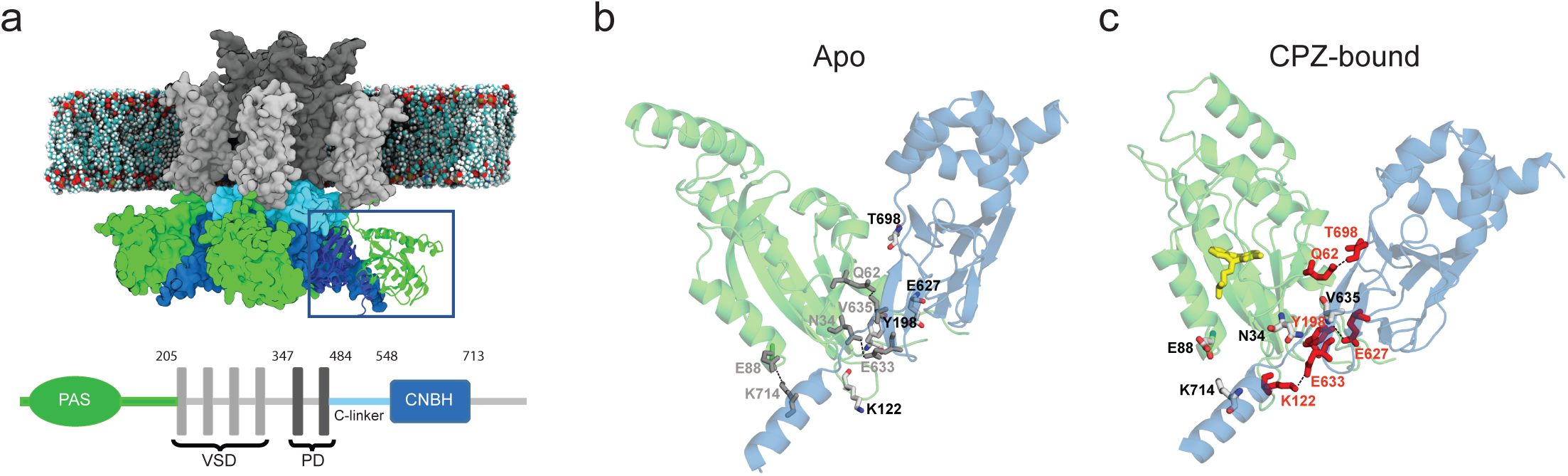
CPZ alters interactions between the PAS and CNBH domains in full-length EAG1 channels. a, An atomistic model of the full-length mEAG1 channel embedded in a membrane composed of POPC lipids. The PAS domain is shown in green, S1-S4 transmembrane helices forming the voltage sensor domain in light grey, S5-S6 transmembrane helices forming the pore domain in dark grey, the C-linker connecting the CNBH domain to the S6 helix in cyan and the CNBH domain in blue. In the bottom, a linear representation of EAG1 channel subunit topology with residue positions at the border of the domains indicated for clarity. The domain color-coding is the same as for the atomistic model shown above. H-bonds and salt-bridges between the interacting PAS and CNBH domains for representative poses in an apo (**b**) and CPZ-bound (**c**) states with the interacting residues labeled and shown in sticks. Residues that form stronger interactions in the apo state are shown in grey in (b) and residues that form stronger interactions in the CPZ-bound state are shown in red in (c). PAS and CNBH domains are show in a semi-transparent representation for clarity.

Analysis of the MD simulations showed that overall the root mean squared deviations (RMSD) in the presence and absence of CPZ, and the root mean squared fluctuations (RMSF) between the apo and CPZ-bound states for the individual domains of the channel were small for the duration of the simulations (Supplementary Fig. 1).

However, comparison of the H-bonds and salt-bridges at the PAS and CNBH domain interface computed in the apo (Fig. 1b) and CPZ-bound (Fig. 1c) states indicated that while some hydrogen bonds and salt-bridges were the same between the two states, the probability of forming or breaking of many bonds changes with CPZ binding (Supplementary Table 1). For instance, interactions between residues N34-E633, Q62- V635 and E88-K714 were weakened after the ligand binding, while interactions between Q14-Y666, Q62-T698, K122-E633 and Y198-E627 were strengthened in the CPZ- bound state. Therefore, the MD simulations indicate that CPZ binding causes a rearrangement of residue interactions at the PAS and CNBH domain interface.

### CPZ strengthens interactions between the isolated PAS and CNBH domains of EAG1 channels

MD simulations suggest that CPZ binding may affect PAS and CNBH domain interaction affinity as some bonds at the PAS and CNBH domain interface are weakening and some are strengthening as a result of CPZ binding. However, experimentally determining the total effect of CPZ binding on the affinity of PAS and CNBH domain interactions in the intact full-length EAG1 channels is not feasible.

Previously, we have shown that surface plasmon resonance (SPR) can be used to determine the affinity of the isolated PAS and CNBH domain interactions in ERG1 channels homologous to EAG1 [31]. Therefore, to experimentally test if CPZ binding can affect interactions between the isolated EAG1 PAS and CNBH domains with SPR, we immobilized the isolated CNBH domains of EAG1 channels on the CM5 sensor chip and applied the isolated PAS domains over the range of concentrations in the absence and presence of CPZ (Fig. 2). The experiments were repeated on three different CM5 chips. The SPR response increased with the increase in the PAS domain concentration in the absence and also in the presence of CPZ. To determine the interaction affinity between the PAS and CNBH domains, the SPR response profiles were fitted with the two-state reaction model, using Biaevaluation software version 1.0, as we have described before [31;32]. The association and dissociation rate constants determined from the fits over the range of the tested PAS domain concentrations for each of the three experiments are listed in the Supplementary Table 2. The rate constants were used to calculate the affinity of the PAS and CNBH domain binding. The averaged PAS and CNBH domain interaction affinity determined in the absence of CPZ was 2.5 ± 0.3 μM. This affinity is slightly higher than the affinity for EAG1 PAS and CNBH domain interactions of 13.2 + 2.3 μM, determined using the fluorescence anisotropy [18]. The difference in the affinities is not surprising as ligand binding affinities determined using different methods frequently differ [33]. Increasing the CPZ concentration increased the PAS and CNBH domain interaction affinity determined with SPR in a concentration dependent manner, from 2.5 ± 0.3 μM in the absence of CPZ to 0.9 ± 0.3 μM in the presence of 50 μM CPZ (Fig. 2, Supplementary Table 2). Thus, the SPR experiments revealed that CPZ strengthened interactions between the isolated PAS and CNBH domains in a concentration-dependent manner.

**Fig. 2.**
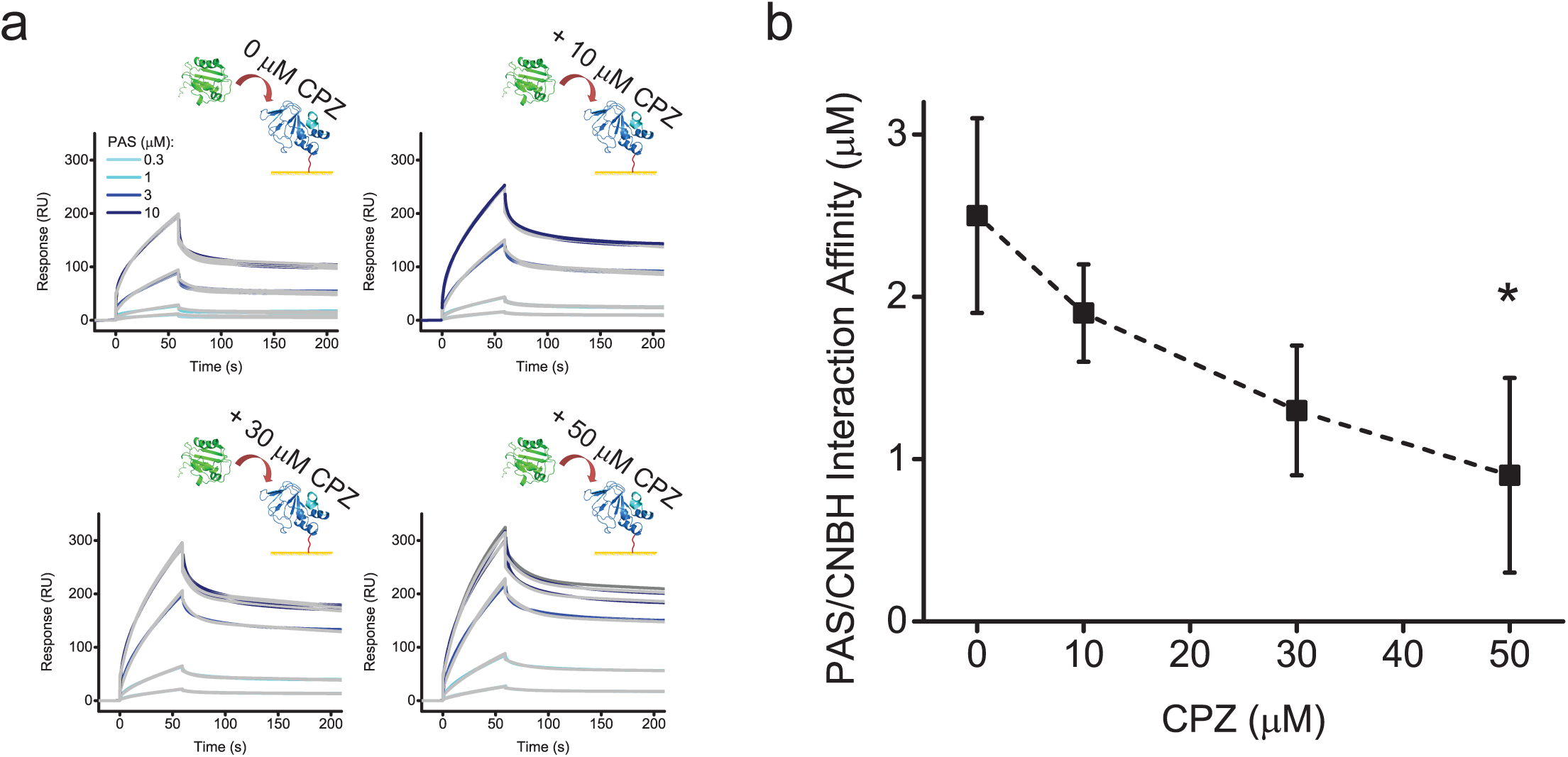
CPZ strengthens interactions between the isolated PAS and CNBH domains. a, SPR sensorgrams for the EAG1 PAS domains applied at the indicated concentrations to the EAG1 CNBH domains immobilized on the CM5 sensor chip in the absence and presence of the indicated CPZ concentrations. Grey lines represent fits of the data with the two-state reaction binding model using the Biaevaluation software. Kd values were 2.9, 1.8, 1.6, and 0.5 µM in the presence of 0, 10, 30 and 50 µM CPZ, respectively. **b**, Plots of the averaged PAS/CNBH domain interaction affinity versus CPZ concentration. The corresponding association and dissociation rate constants for each of the three experiments and the averaged Kd values can be found in the Supplementary Table 2. The data are presented as mean ± SD. * P < 0.05 by ANOVA.

### CPZ inhibits information flow to the pore domain of EAG1 channels

In order to inhibit current flow through EAG1 channels, the conformational changes induced by CPZ binding to the PAS domain have to be communicated to the pore domain via a network of residue-residue interactions. To shed light on the allosteric pathways involved in the transition between the apo and CPZ-bound states, we employed MD network analysis. Network analysis was recently used to investigate the coupling between the VSD and PD in the voltage-gated potassium channel KCNQ1 [34]. For the network analysis, MD trajectories were converted to a residue interaction network, where each node corresponded to an individual amino acid residue within the full-length EAG1 channel. The weights on the edges (connection between nodes) were defined by the spatial proximity and correlated motions of residues in the MD trajectories. The final network encoded all residues (nodes) and interactions (edges) in the channel. The allosteric pathway was measured by calculating the flow of information through the network by defining the PAS domain (binding pocket) as the source of information and the channel pore as the sink. We assumed that in order to inhibit the current flow through EAG1 pore, perturbations of interactions between residues in the PAS domain induced by CPZ binding will have to spread via diffusion in the network to the residues in the pore domain. To identify the key residues and pathways, current (information) flow betweenness analysis was performed to account for all pathways between the source and sink residues [34;35]. Current flow analysis projected onto the structure of the full-length EAG1 channel, with higher information flow (IF) shown in darker colors (dark to light red) with moderate and low IF in lighter colors (orange to yellow), indicated that the CNBH domain region carries most of the information flow to the pore domain, followed by the PAS domain, suggesting that these two domains are allosterically coupled to the pore domain (Fig. 3a-c). The IF for the VSD and PAS domain interface was weaker compared to the PAS and CNBH domain interface (Fig. 3a), suggesting that the allosteric coupling between PAS and VSD is weaker than between PAS and CNBH domains.

**Fig 3.**
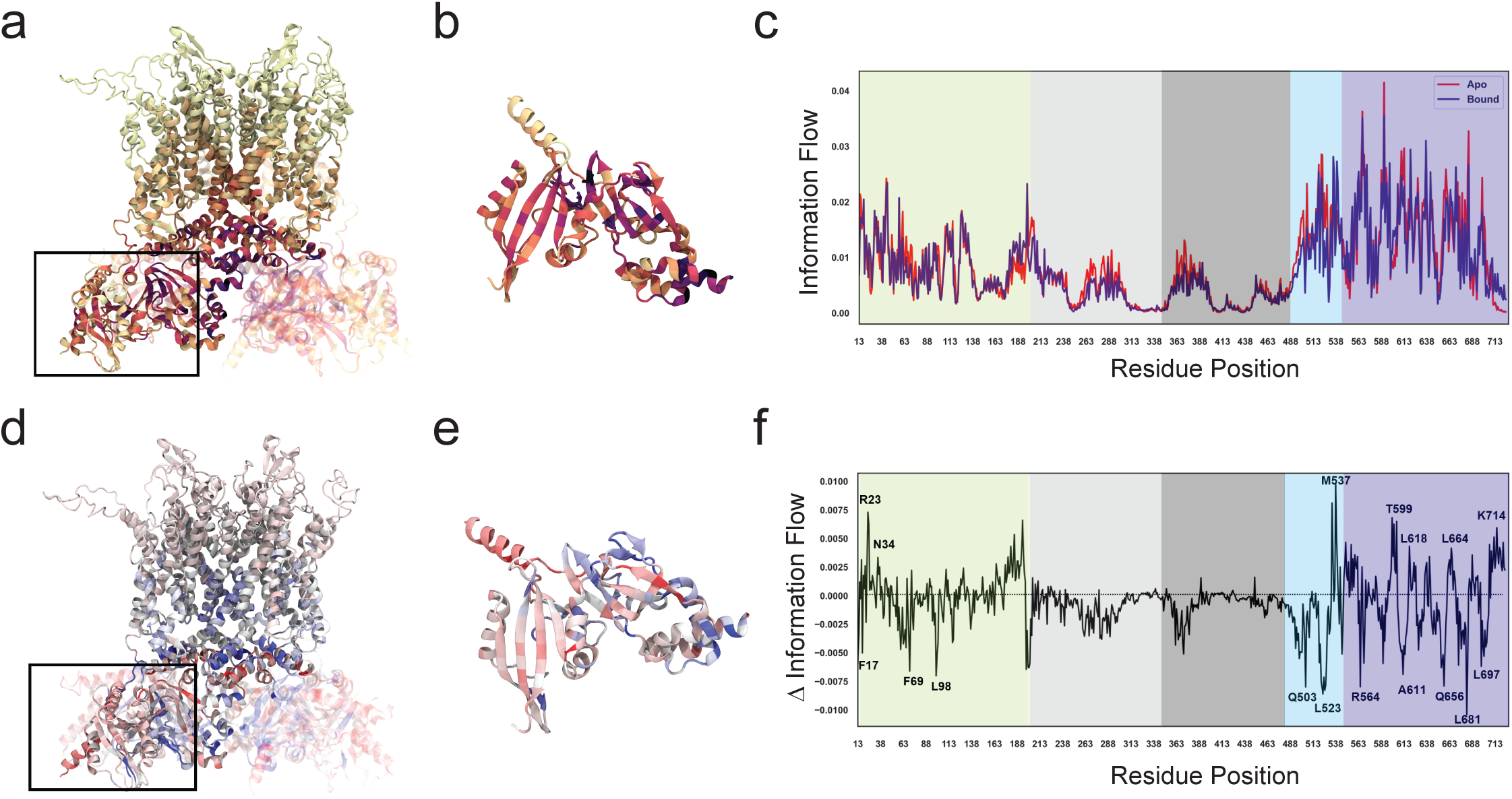
Network analysis of MD simulations indicates that CPZ binding decreases allosteric coupling between the PAS/CNBH domain ring and pore domain. **a**, Information flow through each EAG1 channel residue projected onto the full-length EAG1 structure model generated with MD simulations. Darker colors (dark to light red) have high information flow with moderate and low information flow in lighter colors (orange to yellow), **b**, Enlarged PAS/CNBH domain interaction complex with PAS and CNBH domains from adjacent subunits from the region outlined in **a**. **c**, Information flow profile for the apo (red) and CPZ-bound (blue) EAG1 channels mapped to the amino acid sequence of EAG1 channels. Sequence regions corresponding to the PAS, VSD, PD, C-linker and CNBH are shaded in green, light gray, dark grey, cyan and blue colors, respectively. **d**, Δ information flow, corresponding to the difference between the information flow profiles of apo and CPZ-free states, projected onto the structure of EAG1 channels. Blue color corresponds to the maximal decrease in the information flow between the two states (-0.01) and red to the maximal increase (0.01). **e**, Enlarged PAS/CNBH domain interaction complex with PAS and CNBH domains from adjacent subunits from the region outlined in **d**. **f**, Δ information flow profile mapped to the amino acid sequence of EAG1 channels. Channel domains are shaded as in **c**.

To investigate how CPZ binding to the PAS domain influences the allosteric network, we calculated the difference in IF profiles between the apo and CPZ-bound states of EAG1 channel (ΔIF). The resulting ΔIF projected onto the full-length EAG1 structure revealed overall a lower IF in the PAS and CNBH domain regions and specifically at the PAS-CNBH domain interface in the CPZ-bound state (Fig. 3d-f).

Some of the residues with lower IF in the bound state were F17, F69, L98 on the PAS domain and Q503, L523, R564, A611, Q656, L681, and L697 on the CNBH domain, while residues with higher IF in the bound state were R23, and N34 on the PAS domain and M537, T599, L618, L664 and K714 on CNBH domain (Fig. 3f). Noteworthy, PAS domain residues interacting with the intrinsic ligand, a conserved motif in KCNH channels that occludes CNBH domain cavity [36–38], had lower IF in the CPZ-bound state. Although overall IF decreased for the C-linker/CNBH domain, the distal C-terminal region showed a higher value of IF in the CPZ-bound state. Concurrent decrease in IF for PAS/CNBH domain interface and for the entrance to the pore suggests that the allosteric coupling between the PAS/CNBH domain tetrameric ring and pore domains decreased due to the CPZ binding, leading to the current inhibition. Therefore, the key allosteric pathways are inhibited in the CPZ-bound state.

### CPZ binding to the PAS domain does not affect Cole-Moore shift in EAG1 channels

The network analysis indicated that coupling between the VSD and PAS domain was mostly unchanged due to the CPZ binding to the PAS domain (Fig. 3a). To experimentally test the absence of coupling between the VSD movement and CPZ binding, we used Cole-Moore shift that reflects slow activation of the channel from deep hyperpolarized states [23;39]. Cole-Moore shift is prominent in EAG1 channels and indicates that in order to reach an open state from an extreme hyperpolarized states, VSD of EAG1 channels has to transition through multiple closed states [23;40]. To determine if binding of CPZ to the PAS domain affects the VSD movement through the closed states, we recorded the Cole-Moore shift in the absence and presence of CPZ for currents recorded with the two-electrode voltage-clamp (TEVC) from EAG1 channels expressed in Xenopus *laevis* oocytes. The voltage protocol used for the Cole-Moore shift and elicited currents are illustrated in Supplementary Figure 2a and b. Previously, we have shown that for currents recorded with TEVC, IC50 for EAG1 current inhibition by CPZ is 30 μM [25]. Therefore, the Cole-Moore shift was recorded in the absence (Fig. 4a, black trace) and presence of 30 μM CPZ (Fig. 4a, red trace). The Cole-Moore shift was further analyzed by plotting the averaged normalized currents recorded at 10 ms following the prepulse against the prepulse potentials (Fig. 4b), as in [19], and time to reach 80% of maximal current at the end of the test pulse against the prepulse potential (Fig. 4c), as in [22]. The plots in Fig. 4b were fit with the Boltzmann function to determine V1/2 - the prepulse potential that produces half maximal Cole-Moore shift. CPZ had little effect on the V1/2, indicating that it does not substantially affect VSD movement between the deep hyperpolarized states to the open state.

**Fig. 4.**
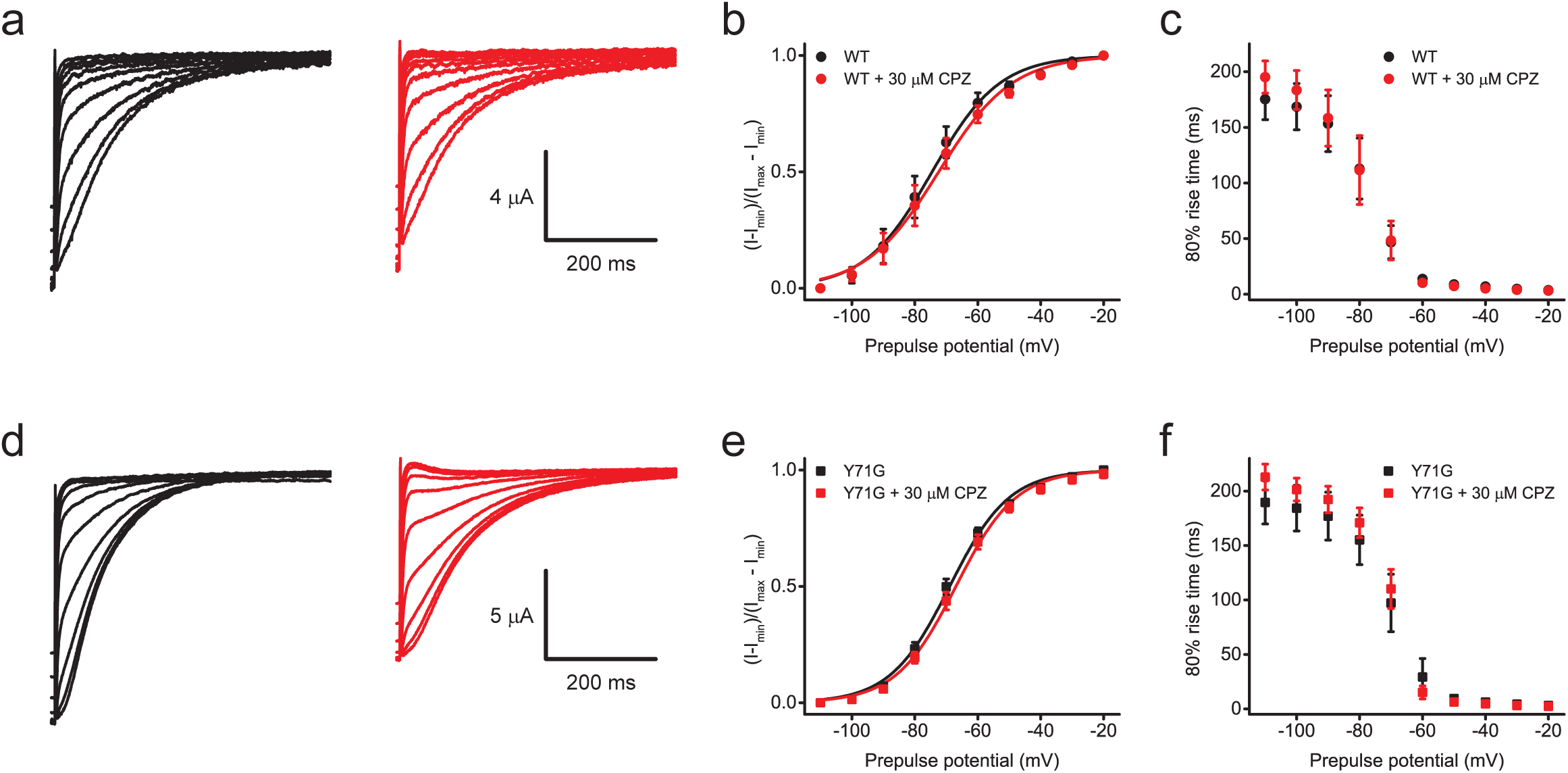
CPZ binding to the PAS domain does not alter Cole-Moore shift in EAG1 channels. Representative currents from WT EAG1 (**a**) and Y71G mutant (**d**) channels recorded at +40 mV following prepulses from -110 mV to -40 mV in the absence (black) and presence (red) of 30 μM CPZ. Plots of the averaged normalized currents for WT EAG1 channels recorded at +40 mV plotted against the prepulse potential in the absence (black symbols) and presence (red symbols) of 30 μM CPZ for WT EAG1 (**b**) and Y71G mutant (**e**) channels. The lines represent fits with the Boltzmann equation with the V1/2 of -74.5 ± 0.7 mV in the absence and -72.4 ± 0.7 in the presence of CPZ for WT channels, and V1/2 of -68.9 ± 0.5 mV in the absence and -66.9 ± 0.5 in the presence of CPZ for Y71G mutant channels. n = 6 for WT channels, and n = 9 for Y71G mutant channels. Plots of the averaged rise time to 80% of the maximal currents at the end of the test pulse plotted against the prepulse potential in the absence (black symbols) and presence (red symbols) of 30 μM CPZ for WT EAG1 (**c**) and Y71G mutant (**f**) channels. n = 6 for WT channels, and n = 4 for Y71G mutant channels. The data are presented as mean ± SD.

We have shown that Y71 in the PAS domain restricts access of CPZ to the PAS domain cavity [26]. Mutating Y71 to a smaller glycine or valine increases EAG1 channel inhibition by CPZ through binding to the PAS domain. Therefore, we tested if Y71G mutation can unmask the effect of CPZ on Cole-Moore shift. Consistent with our previous findings, currents from Y71G mutant EAG1 channels showed higher inhibition by CPZ than the currents from wild-type (WT) channels (Fig. 4d). However, similar to WT channels CPZ did not affect the V1/2 and rise time to 80% of the maximal currents (Fig. 4e and f). Therefore, CPZ binding to the PAS domain does not alter VSD transition from deep hyperpolarized states to the open state.

## DISCUSSION

In this study, we show that binding of small molecule ligand CPZ to the PAS domain of EAG1 channels alters interface between the PAS and CNBH domains without altering the movement of the VSD from deep hyperpolarized to open states. The CPZ-induced changes in the interactions between the PAS and CNBH domains cause a decrease in the coupling between the intracellular domains and channel pore, inhibiting the current through EAG1 channels. Our findings provide the first glimpse into the allosteric pathways involved in EAG1 channel regulation by small molecule ligands targeting their intracellular PAS domains.

It has been suggested that opening of EAG1 channels is mediated by a rotation of the intracellular tetrameric ring formed by the PAS and CNBH domains that leads to a rearrangement of the PD and opening of the channel [19]. Indeed, comparison of the closed and partially open structures of EAG1 channels suggests that the opening of the channel is due to a rotation of the intracellular domains [19]. Moreover, it was proposed that stabilization of the interactions between the PAS, VSD and CNBH domains by calmodulin inhibits currents from EAG1 channels [19]. Further support for the theory that opening of KCNH channels is facilitated by the rearrangements of the PAS and CNBH domains is provided by studies on the EAG-like (ELK) channels based on the combination of patch-clamp fluorometry and incorporation of a fluorescent non-canonical amino acid [41]. Studies on ERG channels using conformational-sensitive antibodies also revealed alterations in the PAS/CNBH tetrameric ring assembly during the channel gating [42] and a more recent report based on the combination of electrophysiology and MD simulations also indicated rearrangements between the intracellular domains and rotation of the tetrameric PAS/CNBH domain ring during the ERG channel gating [43]. Our MD simulations on the full-length EAG1 channels indicate that ligands, such as CPZ, can also alter interface between the PAS and CNBH domains (Fig. 1 b and c). Our computational results are further supported by SPR results that indicate strengthening of interactions between the isolated PAS and CNBH domains in the presence of CPZ (Fig. 2). Although, it is not possible to extrapolate the results for the isolated domains in the context of full-length channels, both MD simulations and SPR results indicate that binding of CPZ to the PAS domain alters the PAS and CNBH domain interface. The CPZ-induced changes in the PAS and CNBH domain interactions could affect the rotation of the tetrameric PAS/CNBH domain ring and the coupling of this rotation to the pore of the channel, inhibiting channel opening in the similar manner as proposed for calmodulin.

The network analysis provides further support for the effect of CPZ via changing the coupling between the tetrameric PAS/CNBH domain ring and the pore of the channel. This analysis indicates that CPZ decreases the information flow between the PAS/CNBH domains and the pore of the channel while having little effect on the information flow between the VSD and PAS domains (Fig. 3d). Since the opening of the channels is thought to be mediated by the rotation of the tetrameric ring, the network analysis results then suggest that the inhibition of EAG1 currents by CPZ is caused by either decoupling or reduced coupling between the tetrameric ring and the pore of the channel while maintaining the network of interactions between the PAS domain and VSD. The analysis of the Cole-Moore shift in the presence of CPZ provides further support that the CPZ binding to the PAS domain does not alter VSD movement, at least from deep hyperpolarized closed states to the open state. Cole-Moore shift is thought to reflect the transition of the VSD between multiple closed states in order to reach an open channel state [23]. In EAG1 channels, the Cole-Moore shift is mediated by the PAS domain via interactions with the VSD and CNBH domains [19-22;40]. Deletion of the PAS and CNBH domains or swapping EAG1 PAS and CNBH domains with the corresponding domains of homologous ERG1 channels removes the Cole-Moore shift [19]. Consistent with this reports, we also did not observe a Cole-Moore shift for EAG1 channels lacking the PAS domain, both in the absence and presence of CPZ (Supplementary Fig. 2c-e). Importantly, CPZ had no effect on the Cole-Moore shift in the WT EAG1 channels and also mutant Y71G EAG1 channels with enhanced CPZ sensitivity (Fig. 4). These results indicate that CPZ does not affect VSD movement between deep hyperpolarized and open states. Therefore, while CPZ alters the PAS/CNBH domain interactions, VSD acts as an anchor for the tetrameric ring potentially contributing to the decreased coupling between the ring rotation and the pore. Our previous studies also indicated that EAG1 channel inhibition by CPZ is mostly voltage independent [25], further suggesting that CPZ binding to the PAS domain does not alter its interactions with the voltage sensor. Combining the results presented in the current study with previously proposed gating rearrangements in EAG1 channels, we propose a model for EAG1 current inhibition by CPZ binding to the PAS domain (Fig. 5). In this model, EAG1 channel opening is mediated by the rotation of the PAS/CNBH domain tetrameric ring facilitated by the VSD movement in response to membrane depolarization [19]. CPZ binding to the PAS domain alters PAS and CNBH domain interactions (potentially stabilizing them) while having little effect on the interactions between the tetrameric ring and VSD. This then decreases the coupling between the tetrameric ring and PD, inhibiting the rotation of the tetrameric ring and favoring the closed state of the channel.

**Fig. 5.**
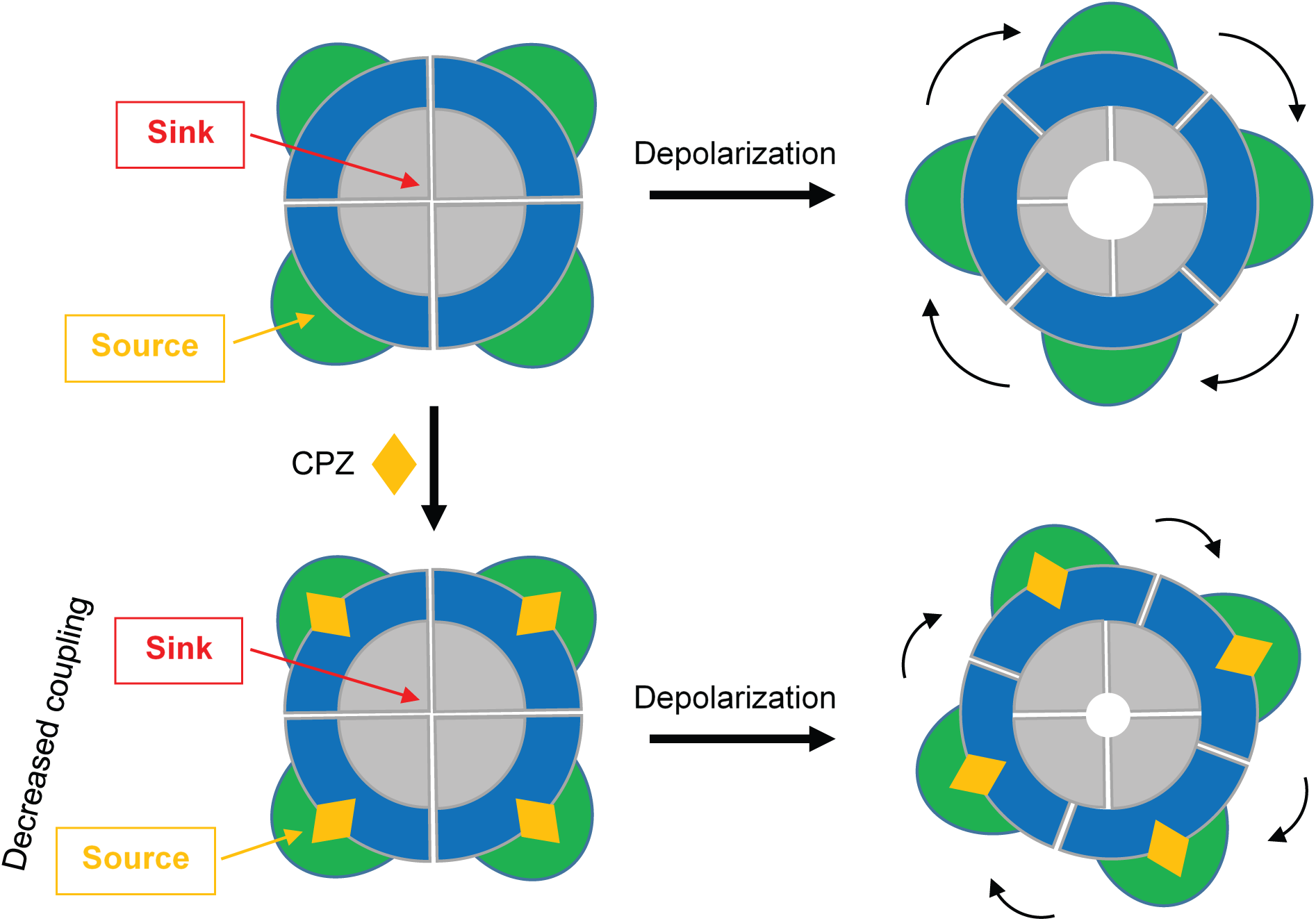
A model of EAG1 current inhibition by CPZ binding to the PAS domain. Depolarization causes a rotation of the intracellular tetrameric ring formed by the PAS and CNBH domains around the pore formed by PDs of the four channel forming subunits. This leads to the opening of the channel pore (top right). PAS domain residues act as a source and PD residues act as a sink. Binding of CPZ, to the PAS domains alters the PAS/CNBH domain interface decreasing the coupling between the source and sink residues (bottom left). This hinders the rotation of the tetrameric ring, inhibiting current from EAG1 channels (bottom right). PAS domains are shown in green, CNBH domains in blue, PD domains in grey, and CPZ in orange.

KCNH channels are the only known ion channels that contain a PAS domain [9;44]. Although distinct in their amino-acid sequence, structurally the PAS domains of KCNH channels share homology with an expansive family of PAS domain-containing proteins in which the PAS domains serve as small molecule binding modules [27;45]. Recent studies firmly established that PAS domains can function as ligand-binding domains in KCNH channels, providing a unique mechanism of current regulation for this family of potassium channels [25;26]. It has been shown that the PAS domains of KCNH channels harbor binding sites for structurally distinct ligands, including tricyclic drugs, such as chlorpromazine and imipramine [25;26] in EAG1 channels, heme [46] in ERG3 channels and undecylenic acid [24] in EAG1 channels. Considering the evolutionary function of PAS domains as ligand-binding modules and their suitability for drug targeting [30], identification and design of additional KCNH PAS domain ligands is imminent. This raises the importance of mechanistic understanding of KCNH channel modulation by the PAS domain ligands. Our study provides the first account of allosteric pathways involved in EAG1 current modulation by the PAS domain small molecule ligands. Most likely similar mechanism applies for EAG1 channel modulation by other tricyclic drugs, including imipramine. Future studies will determine if the pathways of regulation are conserved for other KCNH channels and PAS domain small molecule ligands.

## Author contributions

ZJW and TIB conceived the experiments. ZJW performed electrophysiology experiments. ZJW and TIB analyzed electrophysiology experiments. MG and JBK performed MD simulations and network analysis. XC purified protein for the SPR experiments. PBT performed SPR experiments and data fitting. XC, PBT and TIB analyzed the SPR data. TIB wrote the manuscript with the input from all authors.

## Acknowledgements

This work was supported by grants 1R01CA252969 and R01GM124020 to TIB. SPR sensorgrams were collected and analyzed using the Biacore Molecular Interaction Shared Resource (BMISR) facility at Georgetown University Medical Center. The BMISR is supported by grant P30CA51008. The authors would like to thank Dr. Bernard R. Brooks at the Laboratory of Computational Biology at the National, Heart, Lung and Blood Institute, and the Biowulf high performance Linux cluster at NIH for providing computational resources for this project.

## Methods

### Molecular dynamics simulations of full-length EAG1 channels

For simulating the full-length EAG1 channel in the CPZ-unbound (apo) and CPZ-bound states we first prepared an apo state full-length mouse EAG1 (mEAG1) model using the rat EAG1 (rEAG1) structure determined in the complex with calmodulin (pdb id: 5K7L) [17]. The apo state was prepared in the absence of calmodulin in CHARMM-GUI with 500 POPC lipids per leaflet, AMBER99SB-ILDN parameters for the protein [47] and AMBER parameters for the lipids. The protein-membrane system was solvated with TIP3P water and Na^+^ and Cl^−^ ions were added to a final concentration of 150 mM. To simulate the CPZ-bound state, we aligned the CPZ-bound PAS domains generated from the REST2 docking described in [26] with the PAS domains of each EAG1 subunit in the tetramer and used the coordinates of the CPZ-bound PAS domains for MS simulations of the full-length CPZ-bound EAG1 channels. This was done because in the crystal structure of rEAG1, the Y60 residue (Y71 in mEAG1) blocks the binding of CPZ to the PAS domain cavity and, in order to accommodate CPZ, Y71 swings away from the PAS domain cavity. The system for the CPZ-bound EAG1 was then prepared in the same manner as for the apo state. Both apo and bound states embedded in membrane were equilibrated according to the CHARMM-GUI six-step procedure. During the equilibration phase, the temperature was controlled with Berendsen thermostat and the pressure was maintained at 1bar using a Berendsen barostat. For the production run, we used Nose-Hoover thermostat with coupling constant of 1ps^−1^ to maintain temperature at 310K and Parrinello-Rahman barostat with a compressibility of 4.5×10^−5^ bar^−1^ for the pressure. The simulations ran for 1μs for each apo and CPZ-bound systems.

### Current flow analysis from MD simulations

To analyze allosteric pathways elicited by the CPZ binding to the PAS domain in the full-length EAG1 channels we performed current flow analysis through the channel following the approach laid out by Delemotte’s research group [34;48]. First a continuous contact map was calculated given the distance dij(t) between atoms i and j at time t using truncated Gaussian kernel that is K(dij(t) = 1 if dij(t) ≤ c and otherwise:

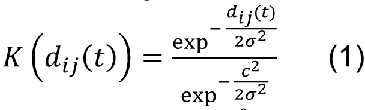

the cutoff c = 4.5 Å was used for the heavy atoms (non-hydrogen atoms) in the simulation. We also used K(dcut) = 10^−5^ for dcut = 0.8 nm leading to σ ≈ 0.138. The final contact map was then averaged over frames. Correlation of residue movements was calculated through mutual information (MI). MI between residues si and sj was estimated based on distances from their equilibrium positions where the position of each residue was defined as the centroid of heavy atoms in each residue. Thus MI is calculated as:

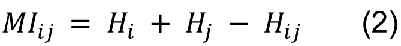

Where Hi is the entropy of residue si defined as:

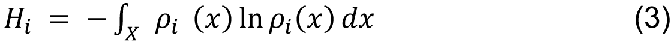

Where the density ρi(x) was estimated using Gaussian mixture model [34]. To increase accuracy, a bootstrapping was performed 10 times and the final MI matrix was averaged over all bootstrap samples. The MI matrix and the semi-binary contact map were used to build the full adjacency matrix Aij = CijIij . This adjacency matrix was used to calculate current (information) flow which measures the flow of information from a set of source (S0) to a set of sink (S1) nodes. The results highlight the nodes that carry the most information from source to sink and give valuable information about allosteric pathways in the protein. For a tetrameric EAG1 protein, the current flow was replicated for each subunit, summed over the structure and then averaged, as was used by the Delemotte’s group to study allostery in KCNQ potassium channels [34]. The source nodes in our analysis were the PAS domain residues (13-138) and the sink nodes were residues at the entrance to the pore of the channel (Q503 on each monomer).

### Protein Expression and Purification

DNA encoding PAS (residues 7-136) and CNBH (residues 552–708) of mEAG1 channels (GI # Q60603) was synthesized by BioBasic (Canada) and subcloned into pETM11 bacterial expression vector containing an N-terminal 6-His affinity tag followed by a tobacco etch virus (TEV) protease cleavage site. The DNA sequences were verified by sequencing (Genewiz). The PAS and CNBH domains were expressed in BL21 (DE3) *Escherichia coli* cells as previously described [25;31]. The cells were grown at 37 °C to an optical density at 600 nm of 0.6-0.8, induced with IPTG at 18 °C overnight and harvested by centrifugation. The cells were resuspended in 150 mM KCl, 1mM TCEP, 1mM ABSF, 2.5 mg/ml DNaseI and 30 mM HEPES, pH 7.5. Cells were lysed with an Emulsiflex-C5 (Avestin). Insoluble protein was separated by centrifugation in 45 Ti rotor at 30,000 rpm for 1hr at 4 °C. The PAS and CNBH domains were purified by Ni^2+^ affinity chromatography using HisTrap HP column (Cytiva) and eluted on a linear gradient to 500 mM imidazole. The protein was further purified on a Superdex 200 Increase 10/300 column (Cytiva) equilibrated with 150 mM KCl, 1mM TCEP, 10% glycerol and 30 mM HEPES, pH 7.5. The protein concentration was determined with Bradford Protein Assay Kit (Pierce).

The purified protein was stored at -80 °C in aliquots and thawed immediately before the experiments. The molecular weight of the PAS and CNBH domains used in the study was verified on Coomassie Blue-stained gels and with mass spectrometry at Proteomics and Metabolomics Core Facility at Georgetown University Medical Center.

### Surface Plasmon Resonance Measurements

All SPR binding experiments were performed on a CM5 chip (Cytiva) at 25 °C using a Biacore T200 Instrument (GE Healthcare). The purified CNBH domains were immobilized on the CM5 chip using a standard amine coupling chemistry in the presence of 10 mM sodium acetate buffer at pH 5.5 as the immobilization buffer (buffer used to directly dissolve ligands), as we described before [31]. HBS-P buffer (150mM NaCl, 10mM HEPES, 0.05% (v/v) surfactant P20, pH 7.4) was used as the immobilization running buffer (buffer that runs in the background during immobilization). Purified PAS domains in the kinetics running buffer (150 mM KCl, 1 mM TCEP, 10% Glycerol, 30 mM HEPES, pH.7.5 and supplemented with 0.05% Tween 20) were injected over the immobilized CNBH domains at 0.3, 1, 3 and 10 μM concentrations in triplicate for 60 s at a flow rate of 50 μl/min (association phase), followed by buffer only injections for 150 s (dissociation phase), in the absence and presence of 10, 30 and 50 μM CPZ dissolved in the kinetics running buffer. Injection of Glycine (pH 2.0) for 15 s was used to regenerate the chip surface. The SPR data were doubly corrected to eliminate the non-specific binding by subtracting the SPR response to the blank (buffer only) injections and also response to the control surface with no immobilized protein.

The buffer for the blank injections contained CPZ at the tested concentrations to make sure that the observed increase in the SPR response for PAS/CNBH interactions is not due to the CPZ binding to the PAS domain but rather due to the effect of CPZ on the PAS/CNBH domain complex formation. CPZ displays non-specific binding to the CM5 chip surface at concentration ≥ 100 μM. Therefore, the effect of CPZ was examined at concentrations ≤ 50 μM. Each experiment was repeated on three different CM5 chips.

To determine the affinity of PAS and CNBH domain interactions in the absence and presence of the indicated CPZ concentrations (Kd), the SPR sensorgrams were fitted with a two-state reaction model based on the assumptions and procedure outlined in our previous publications and used to determine the binding affinity between the PAS and CNBH domains of KCNH2 channels [31;32]. The two-state model describes a biphasic two step conformational change interaction mechanism and is available in the Biaevaluation software version 1.0. Kd is then calculated using the following equation:

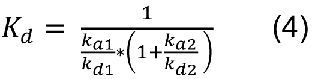

Where ka1 (M^-1^s^-1^) and ka2 (s^-1^) are association rate constants (s^-1^), and kd1 (s^-1^) and kd2 (s^-1^) are dissociation rate constants (s^-1^) for the two-state model.

For all SPR experiments the nonspecific injection signal at the beginning of the association and dissociation phases was excluded from data fitting and figures. Each of the SPR experiments was repeated on three different CM5 chips. The error bars correspond to the S.E. Statistical analysis was performed using Student’s t-tests. P values < 0.05 were considered significant.

### Electrophysiology

The cDNA encoding wild-type (WT) mEAG1 channels (Accession # NP_034730.1) in pGH19 vector was kindly provided by G. Robertson (University of Wisconsin-Madison, Madison, WI). The mutant ΔPAS EAG1 channel in pGH19 with the deletion of the PAS domain residues 2-130 was generated by Bio Basic Inc. (Canada) and verified by DNA sequencing (Genewiz). Mutant EAG1 channels with the amino acid substitutions were generated by Genewiz and verified by DNA sequencing. cRNA was transcribed using the T7 mMessage mMachine kit (Thermo Fisher Scientific). Defolliculated X. laevis oocytes were purchased from Ecocyte Bioscience (Austin, TX) or Xenopus1 (Dexter, MI) and injected with the cRNA using a Nanoinject II oocyte injector (Drummond).

For EAG1 current recordings, oocytes were placed into a handmade chamber containing a bath solution for current recording. The currents were recorded using two-electrode voltage clamp (TEVC) technique with OC-725C amplifier (Warner Instruments) and pClamp11 software (Molecular Devices). The signals were digitized using Digidata 1550 (Molecular Devices). Patch pipettes were pulled from borosilicate glass and had resistances of 0.7–1.5 MΩ. The recording (bath) solution contained 96 mM NaCl, 4 mM KCl, 0.1 mM CaCl2, 1.8 mM MgCl2, and 5 mM HEPES, pH 7.5. Pipette solution contained 3 M KCl. The Cole-Moore shift for WT and mutant EAG1 channels was elicited by applying a series of 7-sec voltage pre-pulses ranging from -110 to +40 mV in 10-mV increments from a holding potential of -80 mV, followed by a 0.5-sec voltage pulse to +40 mV. The currents were not leak-subtracted.

To analyze the Cole-Moore shift we followed the analysis described by Whicher and MacKinnon [19]. Currents at 10 ms following the end of the prepulse were normalized and plotted as a function of the prepulse potential, and the obtained plots were fit with the Boltzmann equation (5):

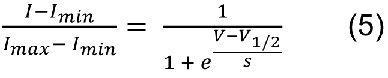

Where 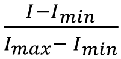 is the normalized current at 10 ms following the prepulse, V1/2 is the voltage that produces half-maximal Cole-Moore shift (mV), and *s* is the slope of the relation (mV).

CPZ was purchased from Alfa Aesar. The CPZ stock and the dilutions were made in the bath solution. For each CPZ concentration, more than five recordings were obtained, each from a different oocyte. The error bars on the figures correspond to the SD. Statistical analysis was performed using One-Way ANOVA. P values < 0.05 were considered significant. The data analysis and fitting of the plots were performed in Clampfit (Molecular Devices) and Origin (Microcal Software, Inc).

---

**Supplementary Fig 1.**
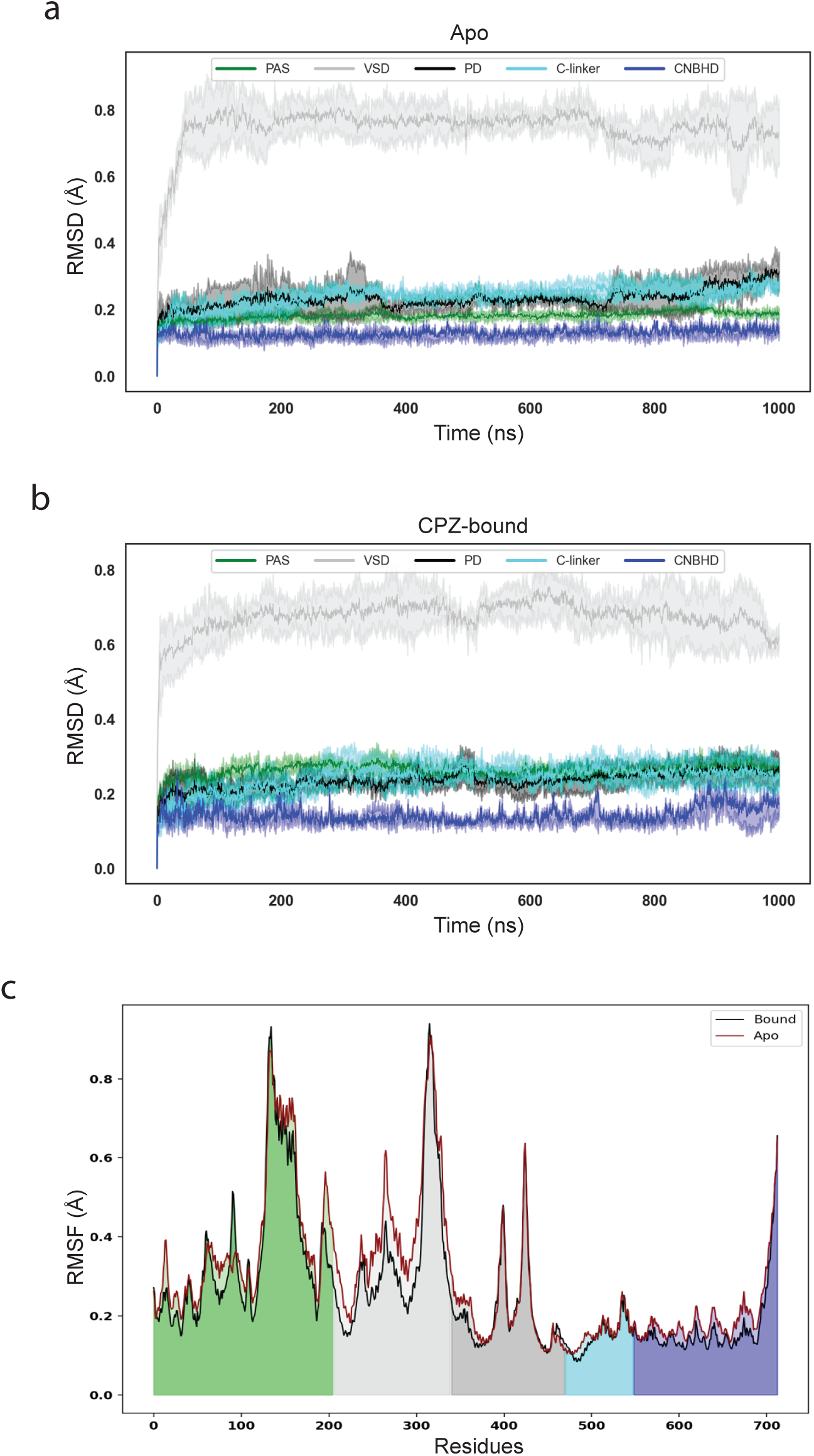
The backbone root-mean-square deviations (RMSD) for different domains of EAG1 channels modeled in apo (**a**) and CPZ-bound states (**b**) for 1 µs MD simulations. The color coding of the domains is the same as in Fig. 1. **c**, The root-mean-square fluctuations (RMSF) between the apo and CPZ-bound states, calculated based on RMSD in (a) and (b), for the indicated residues. Sequence regions corresponding to the PAS, VSD, PD, C-linker and CNBH are shaded in green, light gray, dark grey, cyan and blue colors, respectively.

**Supplementary Fig. 2.**
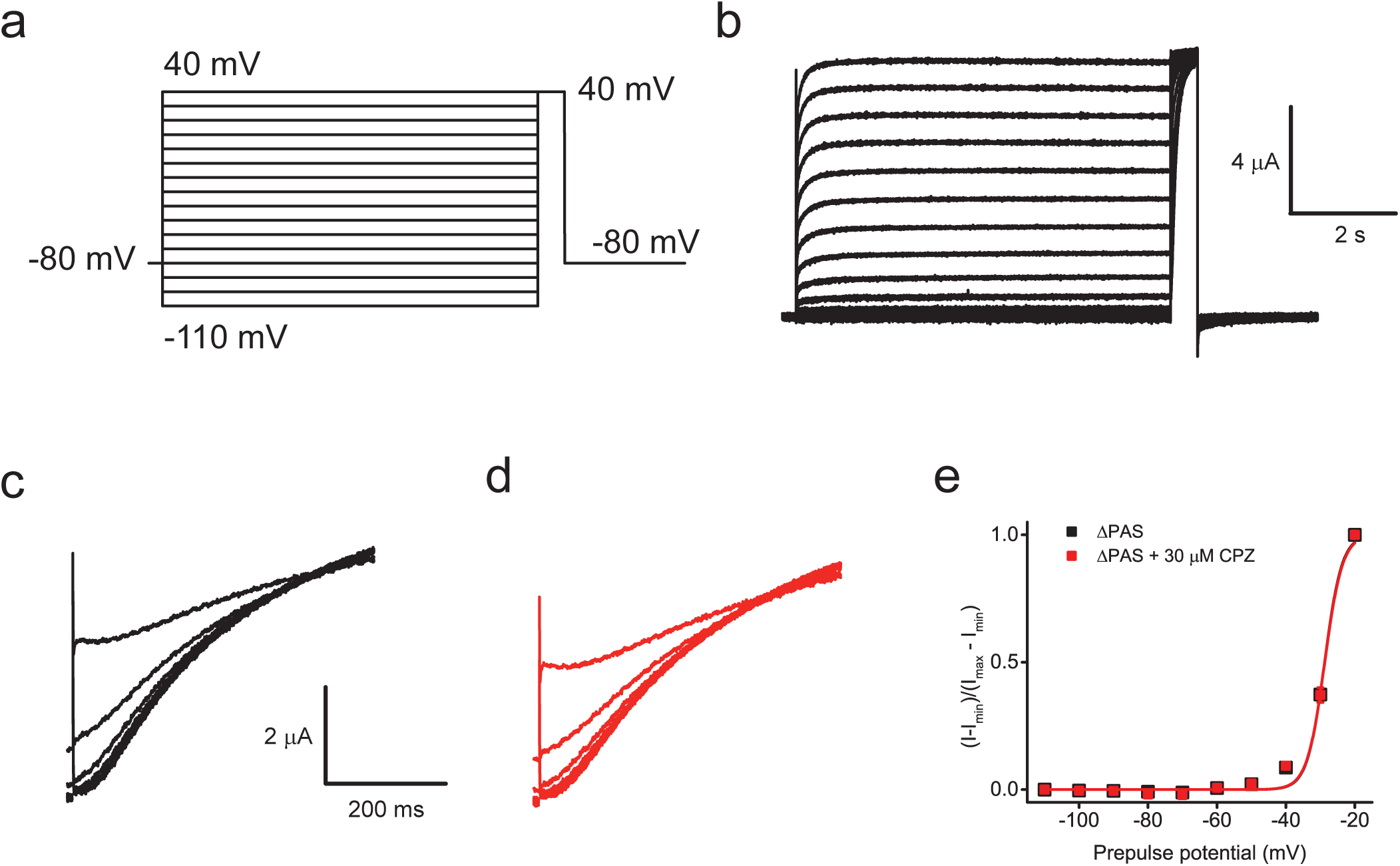
Deletion of the PAS domain abolishes Cole-Moore shift in EAG1 channels. a, Voltage protocol used to elicit Cole-Moore shift in EAG1 channels. **b,** Representative currents from WT EAG1 channels elicited with the voltage protocol shown in (a). Representative currents from mutant EAG1 channels with truncated PAS domain (ΔPAS) recorded at +40 mV following prepulses from -110 mV to -40 mV in the absence (**c**) and presence (**d**) of CPZ applied at 30 μM concentration. The same scale bars in (c) and (d). **e**, Plots of the averaged normalized currents for the ΔPAS EAG1 channels recorded at 10 ms following the prepulse versus the pre-pulse voltage obtained in the absence (black symbols) and presence (red symbols) of 30 μM CPZ. The lines represent fits with the Boltzmann equation with V1/2 of -28.8 ± 0.4 mV in the absence and -28.7 ± 0.5 mV in the presence of CPZ. The black symbols are larger than red symbols to prevent complete overlap of the data in the absence and presence of CPZ. n = 5. The data are presented as mean ± SD.

**Supplementary Table 1.** Probabilities (in percent) of H-bonds and salt-bridges between PAS and CNBH domains in the apo and CPZ-bound states.

| **H-bonds/Salt Bridges** | **Apo (%)** | **CPZ-bound (%)** |
| --- | --- | --- |
| Q14-Y666 | 41 | 45 |
| N15-L664 | 71 | 64 |
| N34-E633 | 89 | 72 |
| N34-V634 | 73 | 74 |
| Q36-I710 | 91 | 92 |
| V38-R708 | 98 | 97 |
| V43-I637 | 43 | 38 |
| Y44-I637 | 26 | 26 |
| R57-D642 | 100 | 100 |
| Q62-V635 | 47 | 24 |
| Q62-T698 | 0 | 21 |
| E88-K714 | 31 | 20 |
| K122-E633 | 2 | 15 |
| Y198-E627 | 70 | 93 |

Residue numbering corresponds to mEAG1 sequence (Accession # NP\_034730.1).
Interactions strengthened by CPZ binding are shown in red, the ones that are weakened in grey, and the ones that are essentially not affected by CPZ binding are shown in black.

**Supplementary Table 2.** Interaction affinity of PAS and CNBH domains in the absence and presence of the indicated CPZ concentrations for SPR experiments performed on three different CM5 sensor chips (n1-n3)

| CPZ ($\mu\text{M}$) | n1 | | | | | n2 | | | | | n3 | | | | | Ave $K_d$ ($\mu\text{M}$) |
| --- | --- | --- | --- | --- | --- | --- | --- | --- | --- | --- | --- | --- | --- | --- | --- | --- |
| | $k_{on1}$ | $k_{off1}$ | $k_{on2}$ | $k_{on2}$ | $K_d$ ($\mu\text{M}$) | $k_{on1}$ | $k_{off1}$ | $k_{on2}$ | $k_{on2}$ | $K_d$ ($\mu\text{M}$) | $k_{on1}$ | $k_{off1}$ | $k_{on2}$ | $k_{on2}$ | $K_d$ ($\mu\text{M}$) | |
| 0 | 1301 | 0.073 | 0.035 | 0.001 | 1.8 | 1395 | 0.112 | 0.035 | 0.001 | 2.7 | 785 | 0.068 | 0.033 | 0.001 | 2.9 | $2.5 \pm 0.6$ |
| 10 | 1247 | 0.051 | 0.036 | 0.002 | 1.7 | 1622 | 0.100 | 0.037 | 0.001 | 2.3 | 1458 | 0.057 | 0.034 | 0.016 | 1.8 | $1.9 \pm 0.3$ |
| 30 | 2050 | 0.002 | 0.010 | 0.543 | 0.8 | 4270 | 0.164 | 0.035 | 0.001 | 1.5 | 1477 | 0.061 | 0.064 | 0.002 | 1.6 | $1.3 \pm 0.4$ |
| 50 | 2193 | 0.002 | 0.001 | 0.775 | 0.7 | 1490 | 0.056 | 0.037 | 0.002 | 1.6 | 1783 | 0.029 | 0.027 | 0.001 | 0.5 | $0.9 \pm 0.6^*$ |

P values were calculated using One-Way ANOVA. $P < 0.05$ was considered statistically significant.

